# Mixed structure- and sequence-based approach for protein graph neural networks with application to antibody developability prediction

**DOI:** 10.1101/2023.06.26.546331

**Authors:** Pranav M. Khade, Michael Maser, Vladimir Gligorijevic, Andrew Watkins

## Abstract

There are hundreds of thousands of known proteins but significantly fewer unique protein folds. Furthermore, proteins often have conserved and even repeating geometric patterns, which can be captured by models of protein structure and function. In this work, we use Delaunay tessellations and ***α***-shapes, which capture these conserved geometric patterns, to define graph adjacency in Graph Convolutional Networks (GCN). We demonstrate the utility of the resulting GCN model on antibody developability prediction. Compared to the frequently used graph adjacencies based on k-nearest-neighbors or a fixed cutoff radius, the Delaunay tessellation and ***α***-shapes better capture residue-specific interactions at a lower computational cost for a given system size. The resulting models achieve state-of-the-art performance on an antibody developability prediction task. Finally, we propose an extension of the model which does not require known or predicted structures but uses an “archetypical” antibody structure to infer likely contacts.

## 1 Introduction

Although living organisms have such phenotypic and genotypic diversity, proteins that are fundamental for almost all biochemical functions still have a limited number of folds [1, 2]. Biochemical environments common to diverse species (e.g., aqueous cytosol; hydrophobic cell membranes) exert a common influence and make certain protein folds more favorable than others [3, 4]. The relative disposition of amino acids in these common folds generates protein variants with diverse properties. Patterns in these folds, if captured accurately, can explain a great deal of variation in protein properties using computational models. Efficient modeling of these properties can be particularly useful for developing novel proteins including therapeutics.

Therapeutic proteins like antibodies must exhibit acceptable biophysical properties to be successful drugs, from low viscosity and low aggregation to high thermal and chemical stability. A handful of these properties are summarized by the concept of “developability”; i.e., whether a molecule could likely be developed into a successful therapeutic. Computational predictors of developability properties such as the Developability Index[5] use traits of antibody sequence and structure to gauge developability. In this study, we develop Graph Neural Network (GNN) models capable of emulating this predictor given a crystal structure or an Immunebuilder-predicted structure[6].

GNNs are rapidly becoming a powerful tool for several applications [7, 8] including those related to proteins[9–13]. GCNs are a class of GNNs that use a graph convolution operation to process node features based on their connectivity and accumulate information used in a predictive model. Semantically meaningless edges at best require unnecessary computation [14] and at worst increase the risk of overfitting. Worse, in extremely deep networks, excessive edge density can lead to a form of information loss called “over-smoothing”[15], making selection of a limited subset of edges even more important.

There are many possible adjacency definitions, particularly for large molecules like proteins with crucial information in their 3D conformations, requiring noncovalent edges. Noncovalent adjacency often uses a radial cutoff or a nearest neighbor count to choose which noncovalent edges to include [16]. These methods can introduce extraneous edges and require tuning a parameter (the radial threshold or neighbor count). Therefore, we explore an alternative to these adjacency schemes using the Delaunay tessellation.

Delaunay tessellations (DTs) [17] have simple rules. Given a set of points in three dimensions *P*, a DT is a set of four points that form a circumsphere that encloses no other points in *P*. A set of all the DTs for the given points is called a ‘Convex Hull’ of these points, and if we select a specific DT based on the radius of the circumsphere formed by a tessellation, then a set of such tessellations is called an *α*-Shape (AS)[18]. These tessellations have already been applied to protein structures[19] to predict protein hinges[20] and model protein packing, to predict protein stability via four-body statistical potentials,[21] and to design elastic network models whose fluctuations align well with experimental B-factors[22]. DT-defined graphs have even been investigated for quantum chemistry applications[23], where they specifically outperform KNN based schemes for function interpolation. Because the DT omits edges with intervening nodes, even a coarse-grained graph with DT-based adjacency will only include edges that could be relevant to protein packing. Even without a threshold parameter DT will still use far fewer edges than a 15 Å cutoff model, and is guaranteed to produce a connected graph. A finite threshold parameter, as in AS models, may be used to reduce the number of edges further. In this study, we use these schemes to train a binary classifier of developability index[5] based on antibody structure, encoding the antibody sequence for node features using either Kidera factors (*H*_*KF*_)[24] or one hot encoding (*H*_*OHE*_). This approach is robust to using predicted antibody structures instead of crystal structures, or to an ‘archetype’ based model that uses contact probabilities for edge weights. The overall schema of the experiment is shown in Figure 2. Our results suggest that DT and AS based adjacency schemes produce graphs sufficient for accurate antibody property prediction[25–27].

**Fig. 1.**
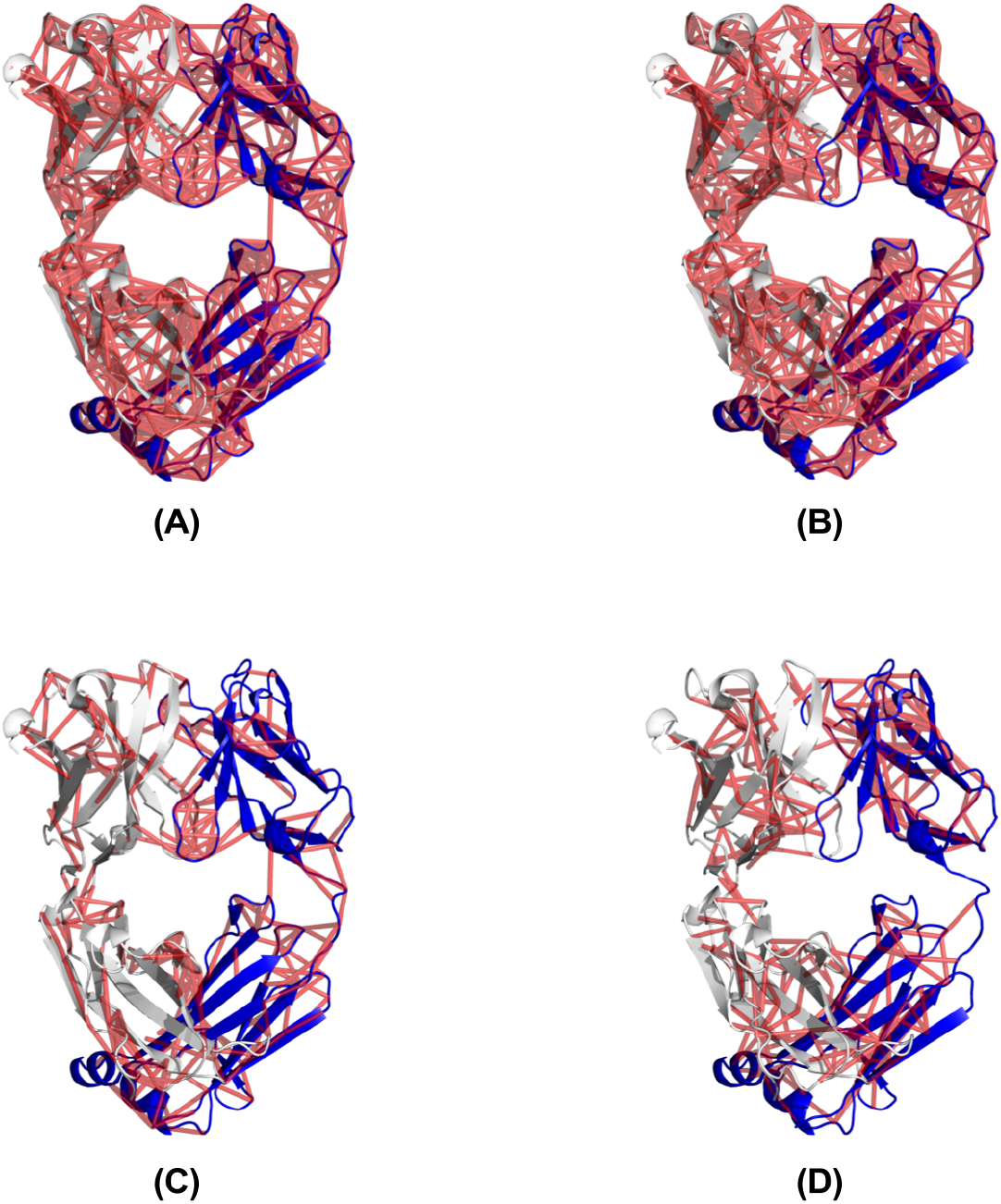
Different adjacency matrices with a comparable number of edges (supplementary table 4) visualized on the Pertuzumab crystal structure (PDB ID: 4LLU). (A) Edges with KNN 6 adjacency with a total of 1,494 edges (B) Edges with DT (*α* = 4 Å) adjacency with a total of 1,516 edges (C) Total 291 edges highlighted that are present in KNN 6 but not in DT (*α* = 4 Å) (D) Total 313 edges in DT (*α* = 4 Å) but not in KNN 6. The number of edges for this structure with all other adjacency schemes is listed in Table S5.

**Fig. 2.**
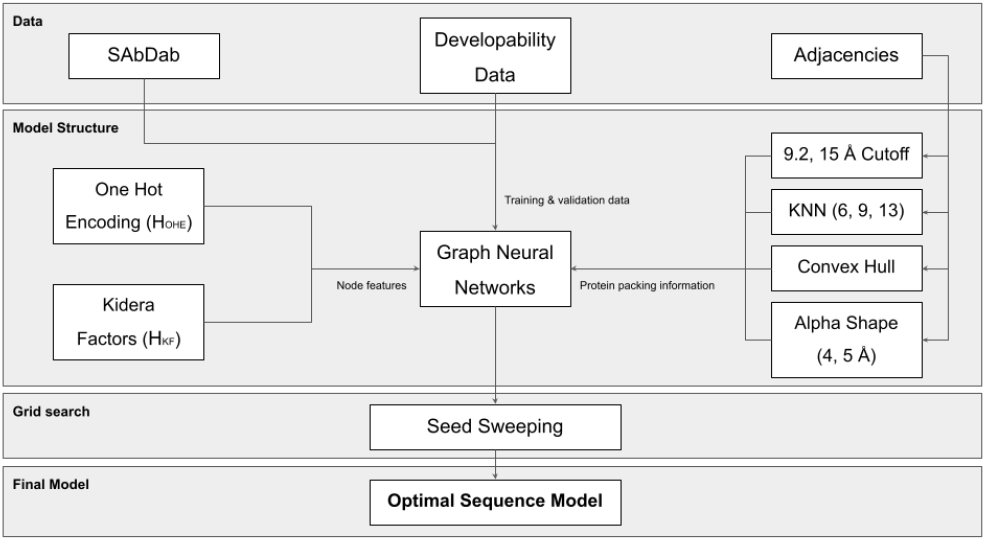
The pipeline architecture is used to build the models in this experiment.

The results for the model employing predicted structures using ImmuneBuilder are noted in Table S4; we do not see a significant change in the performance of the model, which further highlights the robustness of this model architecture. The best model using the antibody archetype (and therefore no structural information specific to the antibody in question) has an F1 score of 0.723 (*σ*:0.04073); worse than the best structure-based models but still superior to the benchmark sequence-based model (Table 1).

**Table 1.** Model performances (15 seeds each). Note: CO means Cutoff

| Adjacency | Max F1 | F1 | Accuracy |
| --- | --- | --- | --- |
| | $H_{OHE}$ | | |
| 15 Å CO | 0.772 | $0.724 \pm 0.039$ | $0.865 \pm 0.025$ |
| 9.2 Å CO | 0.766 | $0.726 \pm 0.032$ | $0.869 \pm 0.024$ |
| KNN 13 | 0.789 | $0.732 \pm 0.036$ | $0.881 \pm 0.02$ |
| KNN 9 | 0.801 | $0.742 \pm 0.041$ | $0.867 \pm 0.021$ |
| KNN 6 | 0.794 | $0.706 \pm 0.052$ | $0.877 \pm 0.024$ |
| DT | 0.768 | $0.717 \pm 0.032$ | $0.861 \pm 0.016$ |
| DT 5 | 0.759 | $0.702 \pm 0.045$ | $0.882 \pm 0.023$ |
| DT 4 | 0.786 | $0.71 \pm 0.036$ | $0.871 \pm 0.021$ |
| | $H_{KF}$ | | |
| 15 Å CO | 0.774 | $0.714 \pm 0.032$ | $0.855 \pm 0.02$ |
| 9.2 Å CO | 0.764 | $0.71 \pm 0.037$ | $0.875 \pm 0.016$ |
| KNN 13 | 0.766 | $0.72 \pm 0.037$ | $0.866 \pm 0.019$ |
| KNN 9 | 0.781 | $0.73 \pm 0.033$ | $0.879 \pm 0.02$ |
| KNN 6 | 0.755 | $0.702 \pm 0.04$ | $0.878 \pm 0.018$ |
| DT | 0.776 | $0.684 \pm 0.062$ | $0.865 \pm 0.024$ |
| DT 5 | 0.79 | $0.712 \pm 0.043$ | $0.884 \pm 0.022$ |
| DT 4 | 0.771 | $0.707 \pm 0.033$ | $0.882 \pm 0.022$ |
| Archetype | 0.723 | $0.6705 \pm 0.04$ | $0.860 \pm 0.022$ |
| Benchmark | NA | $\mu : 0.52 - 0.65$ | NA |

## 2 Materials and Methods

### 2.1 Software

PACKMAN-Molecule [28] and the Geometry module are used to obtain the adjacency matrix representation for the GNN (see the following sections for details).

### 2.2 Datasets

Following Chen et al. [29] we use a subset of SAbDab [30] for the developability model that was derived using BIOVIA’s Pipeline Pilot [31]. In that work, they calculate Developability Index (DI) [5] from the full-length antibody’s net charge and the spatial aggregation propensity of the CDR region. This dataset includes a total of 2409 examples of antibody sequences with corresponding DI. The authors consider a classification task where the lowest 20%, or 482, antibodies by DI are given positive (i.e., highly developable) labels.

There are multiple instances of structures for particular antibody sequences (mainly multiple chains of the same crystal structures). However, only one structure per sequence is selected to prevent test set contamination in cross-validation. Training and validation datasets are split randomly with a 4:1 (80:20) proportion. After splitting, we have set the training batch size as 32 and the validation batch size as 8 following the low validation-low learning rate strategy [32].

We have tested all the adjacencies listed in the first column of Table S1. The thresholds for cutoff, KNN, and AS are selected to keep the average number of edges similar to enable fair comparisons across adjacency schemes. The average number of edges for each adjacency type is noted in Table S1, and to highlight the differences between the two adjacency schemes, Figure 1 shows the edges that are found uniquely in KNN 6 and DT *α* = 4.

### 2.3 Kidera factors as node features

Kidera factors (KF) are 188 physical properties of amino acids which were reduced to 10 properties (Table S2 & S3) with several layers of statistical methods by Kidera and the team [24]. In the last half-decade, the KF has been used in several immunological analyses including Adaptive Immune Receptor Repertoire analysis [33], antigen binding poses of nanobodies (single-domain antibody) re-ranking [34], associations between T cell receptor sequence and gene expression [35, 36], immune repertoire comparison of risk cohorts and patients with COVID-19 with healthy individuals [37], differentiating between polyreactive and non-polyreactive antibodies [38], TCR-peptide binding [39, 40], MHC binding [41], HLA pocket featurization [42] and repertoire analysis [43–49]. In each of these studies, the amino acid alphabet is encoded by KF to build predictive models. Therefore, we also use the same features to represent amino acid nodes in our GNNs/GCNs and compare to one-hot encoding.

### 2.4 Use of antibody archetype

We further extend this model to predict developability index without requiring a pre-existing crystal structure via two strategies. First, we used ImmuneBuilder [6] to predict antibody structures from sequence, then trained a structure-based model to predict developability index. Second, we use prior antibody structures to produce an antibody ‘archetype’ graph with weighted edges. Using Chothia-numbered structures from SAbDab, we build graphs for each structure and include all edges present at least once in the dataset. If an edge between two Chothia-numbered residues is observed in 70% of the structures in the database, then we assign it a weight of 0.7; this weighted adjacency matrix then may be used directly for any antibody sequence. We calculated the archetype graph with DT 4, KNN 6, and Cutoff 9.2 Å (choice as per the same expectation explained in the introduction); we found that DT 4 has 22,396, KNN 6 has 44,529 and Cutoff 9.2 Å has 69,816 unique edges, highlighting the efficiency with which DT 4 adjacency captures protein structural information.

### 2.5 Model Architecture

We are using PyTorch geometric [50] GCN layers from the Kipf et. al. study [7]. Here the hidden features *H* for the *l* + 1^*th*^ convolution layer for a protein with a number of residues R are calculated in the following manner:

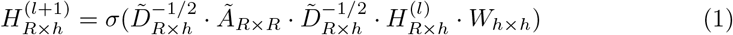

where *A* is an adjacency matrix, *W* is a trainable weight matrix (linear layer), *D* is the marginal sum over A, i.e., 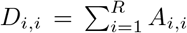, *h* is the dimension of layer *l* + 1, and *σ* is the ReLU activation function. The initial feature matrix of a protein of *R* residues with 10 Kidera Factors each is thus represented as 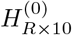.

In our experiment to test different adjacency schemes on a binary problem of developability, we have kept the learning rate of 0.0001, and epoch 600 constant with early stopping upon 60 consecutive epochs of not improving the loss term. We have tested the model with three hidden convolutional layers of size 100 each (100-100-100). Each hidden layer was followed by a Rectified Linear Unit (ReLU) activation function. In the final layer, we have used the global mean pool to return batch-wise graph-level learned output vectors by averaging (mean) node features across the node feature dimension. In each training and validation step, we use a bin with the highest value (torch.argmax) as a predicted class for the binary classification problem. In order to have a statistically significant result, we have run each adjacency and node feature model with different seeds (listed in Table S6) to obtain an average and statistically robust result.

All models herein were trained on a single NVIDIA Quadro P6000 GPU.

## 3 Results

### 3.1 Binary Developability Prediction

Chen et. al. used learned protein embeddings [51] of antibody sequences to train developability predictors [29], where mean validation F1 score with 10-fold cross-validation was mainly considered as a metric for the performance. Support vector machines with the specific combination of physicochemical and learned embedding features were the best performing. Their best models for the test dataset achieved mean F1 scores of 0.52–0.65, which we will use as a baseline for our results.

We have trained several models with different adjacencies and initializations as described in section 2.5. The validation scores of these models are shown in Table 1. Visualizations of results for all the models having node features as Kidera factors (*H*_*KF*_) can be seen in Figures S1 and S3, and those with one-hot encoding (*H*_*OHE*_) in Figures S2 and S4. As can be seen in the results, the average F1 scores for all the models herein are substantially higher than those reported previously. The performance of the best models amongst the seeds are noted as **Max F1**.

The best validation F1 score performance noted by the sweep for *H*_*KF*_ was 0.7898 using the DT 5 model, whose accuracy was 0.8842. For the *H*_*OHE*_ model, the best F1 score performance noted is 0.801 with accuracy 0.8697, achieved by the KNN 9 model.

We also trained and validated the same architecture and *H*_*KF*_ using antibody structures predicted by ImmuneBuilder. The results, noted in Table S4, are very similar to those above, highlighting the robustness of our approach. We hypothesize that this may be because both models are trained on coarse-grained structures (*C*_*α*_ atoms) and may therefore be relatively insensitive to small coordinate changes. For all models, we see a gradual decrease in the loss over the validation steps as seen in Figure S7. For the archetype model, we use the Kidera factors and DT 5 scheme, our best performing *H*_*KF*_ -based model above (DT 5 has many fewer edges; still similar performance as seen in section 2.4). The best F1 score for the archetype-based model is 0.723 (*σ*:0.04073); better than the benchmarked sequence-based model (Table 1). This intermediate performance level is compelling: as one might expect, a graph based on the specific structure of a particular antibody outperforms a ‘generic’ graph based on an archetypical antibody structure, but this archetype model nonetheless outperforms the sequence based prior art, suggesting the DT graph structure provides important biophysical insight.

### 3.2 GPU consumption

GPU consumption is correlated with the number of edges[14]; 15 Å cutoff-based adjacency consumes the highest GPU utilization for a longer period of time and DT (*α* =4) and KNN 6 have the lowest GPU consumption (Figure S9, S10).

Because we chose cutoffs for final model comparisons that kept the number of edges was comparable across adjacency schemes, GPU consumption was similar for those final models. But the number of edges in the antibody ‘archetype’ models varied sharply, even when using cutoffs with comparable edge count for individual structures. That suggests that the edge assignments for DT (*α* = 4), with the fewest total edges (2.4), were best conserved across all the structures summarized into the archetype model. This could suggest that the DT model limits spurious edges that appear transiently in some structures.

## 4 Discussion

Each adjacency scheme for the structure-based GNN model studied used comparable edge count and performed comparably on DI predictions. As to node features, although the performances of both *H*_*KF*_ and *H*_*OHE*_ are not significantly different, *H*_*KF*_ uses ten biophysically meaningful and interpretable features per node, rather than twenty one-hot dimensions. The success of Kidera factors with this model highlights the importance of recalculating Kidera factors, or other amino-acid specific descriptor sets, based on high-quality crystal structures available today. Imposing biologically meaningful priors through edge adjacency and node feature choice helps to ensure interpretability and limit risks from overfitting.

The novel ‘archetype’ approach demonstrates how we can further simplify modeling by circumventing the structure prediction process, using prior structures to encode likely contacts between residues of a novel sequence. Archetype models may extend to property prediction challenges that are limited to individual protein families like immunoglobulins.

Coarse-grained DT GNN models may be applied to predict other antibody properties such as affinity, thermostability, and viscosity, or indeed to other biomolecular property prediction datasets with structural data or accurate structure prediction methods available; in particular, the choice of coarse-graining helps to ensure limited dependence on small structural details.

## 5 Conclusion

We highlight how simple, coarse-grained GNN models can capture a critical aspect of proteins to predict developability. Further, we tested several adjacency schemes and two types of node features to represent protein structure to test its effect on the predictions. It was found that graph formulations of 3D structures based on Delaunay tesselations achieve accuracies near those of traditional radial graphs (e.g., KNN) while encoding far fewer edges. Using our approach, we consistently demonstrate mean F1 scores of 0.70–0.75, substantially improving over prior art. and based on that information, we developed an ‘archetype’ model that demonstrates how we can further simplify the models by circumventing the structure prediction process. Future studies will explore deeper the pruning of potentially spurious edges in moving from KNN-to Delaunay-based GNNs and its effects on model robustness and interpretability.

