## Supplementary Material for "Mixed structure- and sequence-based approach for protein graph neural networks with application to antibody developability prediction"

### Supplementary Tables

**Table S1** The adjacencies are compared with each other. These threshold schemes between different adjacency types are selected to make sure that the adjacency schemes being tested against one another have a comparable number of edges on average.

| Compared Adjacencies | Median number of edges |
|---|---|
| DT 4 and KNN 6 | 1365.137, 1385.487 |
| DT 5 and KNN 9 | 2039.195, 2075.142 |
| DT inf, KNN 13 and Cutoff 9.2 | 2878.585, 2976.288, 2830.476 |

**Table S2** Kidera factors that are used as node features

| Amino Acid | Code | KF1 | KF2 | KF3 | KF4 | KF5 | KF6 | KF7 | KF8 | KF9 | KF10 |
|---|---|---|---|---|---|---|---|---|---|---|---|
| Alanine | ALA | -1.56 | -1.67 | -0.97 | -0.27 | -0.93 | -0.78 | -0.2 | -0.08 | 0.21 | -0.48 |
| Arginine | ARG | 0.22 | 1.27 | 1.37 | 1.87 | -1.7 | 0.46 | 0.92 | -0.39 | 0.23 | 0.93 |
| Asparagine | ASN | 1.14 | -0.07 | -0.12 | 0.81 | 0.18 | 0.37 | -0.09 | 1.23 | 1.1 | -1.73 |
| Aspartic Acid | ASP | 0.58 | -0.22 | -1.58 | 0.81 | -0.92 | 0.15 | -1.52 | 0.47 | 0.76 | 0.7 |
| Cysteine | CYS | 0.12 | -0.89 | 0.45 | -1.05 | -0.71 | 2.41 | 1.52 | -0.69 | 1.13 | 1.1 |
| Glutamine | GLN | -0.47 | 0.24 | 0.07 | 1.1 | 1.1 | 0.59 | 0.84 | -0.71 | -0.03 | -2.33 |
| Glutamic Acid | GLU | -1.45 | 0.19 | -1.61 | 1.17 | -1.31 | 0.4 | 0.04 | 0.38 | -0.35 | -0.12 |
| Glycine | GLY | 1.46 | -1.96 | -0.23 | -0.16 | 0.1 | -0.11 | 1.32 | 2.36 | -1.66 | 0.46 |
| Histidine | HIS | -0.41 | 0.52 | -0.28 | 0.28 | 1.61 | 1.01 | -1.85 | 0.47 | 1.13 | 1.63 |
| Isoleucine | ILE | -0.73 | -0.16 | 1.79 | -0.77 | -0.54 | 0.03 | -0.83 | 0.51 | 0.66 | -1.78 |
| Leucine | LEU | -1.04 | 0 | -0.24 | -1.1 | -0.55 | -2.05 | 0.96 | -0.76 | 0.45 | 0.93 |
| Lysine | LYS | -0.34 | 0.82 | -0.23 | 1.7 | 1.54 | -1.62 | 1.15 | -0.08 | -0.48 | 0.6 |
| Methionine | MET | -1.4 | 0.18 | -0.42 | -0.73 | 2 | 1.52 | 0.26 | 0.11 | -1.27 | 0.27 |
| Phenylalanine | PHE | -0.21 | 0.98 | -0.36 | -1.43 | 0.22 | -0.81 | 0.67 | 1.1 | 1.71 | -0.44 |
| Proline | PRO | 2.06 | -0.33 | -1.15 | -0.75 | 0.88 | -0.45 | 0.3 | -2.3 | 0.74 | -0.28 |
| Serine | SER | 0.81 | -1.08 | 0.16 | 0.42 | -0.21 | -0.43 | -1.89 | -1.15 | -0.97 | -0.23 |
| Threonine | THR | 0.26 | -0.7 | 1.21 | 0.63 | -0.1 | 0.21 | 0.24 | -1.15 | -0.56 | 0.19 |
| Tryptophan | TRP | 0.3 | 2.1 | -0.72 | -1.57 | -1.16 | 0.57 | -0.48 | -0.4 | -2.3 | -0.6 |
| Tyrosine | TYR | 1.38 | 1.48 | 0.8 | -0.56 | 0 | -0.68 | -0.31 | 1.03 | -0.05 | 0.53 |
| Valine | VAL | -0.74 | -0.71 | 2.04 | -0.4 | 0.5 | -0.81 | -1.07 | 0.06 | -0.46 | 0.65 |
| ASN/ ASP | ASX | 0.86 | -0.145 | -0.85 | 0.81 | -0.37 | 0.26 | -0.805 | 0.85 | 0.93 | -0.515 |
| GLN/ GLU | GLX | -0.96 | 0.215 | -0.77 | 1.135 | -0.105 | 0.495 | 0.44 | -0.165 | -0.19 | -1.225 |
| LEU/ ILE | XLE | -0.885 | -0.08 | 0.775 | -0.935 | -0.545 | -1.01 | 0.065 | -0.125 | 0.555 | -0.425 |
| Unknown | XAA | 0 | 0 | 0 | 0 | 0 | 0 | 0 | 0 | 0 | 0 |

**Table S3** Kidera factors and their descriptions.

| Kidera Factor | Description |
|---|---|
| KF1 | Helix/bend preference |
| KF2 | Side-chain size |
| KF3 | Extended structure preference |
| KF4 | Hydrophobicity |
| KF5 | Double-bend preference |
| KF6 | Partial specific volume |
| KF7 | Flat extended preference |
| KF8 | Occurrence in alpha region |
| KF9 | pK-C |
| KF10 | Surrounding hydrophobicity |

**Table S4** $H_{KF}$ model performance for the predicted structures with 14 seeds. We have not built $H_{OHE}$ for the predicted model to avoid redundancy. This model has similar performance to crystal structure-based model.

| Adjacency | F1 | Accuracy | Loss |
|---|---|---|---|
| | **HKF predicted structures** | | |
| KNN 13 | 0.758 | 0.86 | 0.356 |
| KNN 9 | 0.761 | 0.852 | 0.367 |
| KNN 6 | 0.732 | 0.869 | 0.336 |
| DT inf | 0.733 | 0.828 | 0.391 |
| DT 5 | 0.754 | 0.852 | 0.349 |
| DT 4 | 0.761 | 0.864 | 0.351 |
| Cutoff 9.2 | 0.742 | 0.837 | 0.37 |
| Cutoff 15 | 0.722 | 0.856 | 0.374 |

**Table S5** Reduction in the number of edges for the Pertuzumab (PDB ID: 4LLU) for the $C_\alpha$ atoms.

| Edge Type | Number of Edges |
|---|---|
| 15 Å cutoff | 11438 |
| 9 Å cutoff | 3040 |
| KNN (K=6) | 1494 |
| KNN (K=9) | 2235 |
| KNN (K=13) | 3186 |
| DT ($\alpha$ = 4) | 1516 |
| DT ($\alpha$ = 5) | 2266 |
| DT ($\alpha$ = inf) | 3086 |

**Table S6** The seeds used for the sweeps for the result reproducibility.

| Sweep | Seeds |
|---|---|
| $H_{KF}$ | 239, 1223, 2342, 5290, 23, 1235, 7456, 234, 9865, 33, 235, 11, 985, 1260, 5 |
| $H_{OHE}$ | 239,1223,2342,5290,23,1235,7456,234,9865,33,235,11,985,1260,5 |
| $H_{KF}$ predicted structures | 239, 1223, 2342, 5290, 234, 1235, 7456, 9865, 33, 235, 11, 985, 1260, 5 |
| $H_{KF}$ antibody archetype sequence | 239, 1223, 2342, 5290, 234, 1235, 7456, 9865, 33, 235, 11, 985, 1260, 5 |

### Supplementary Figures

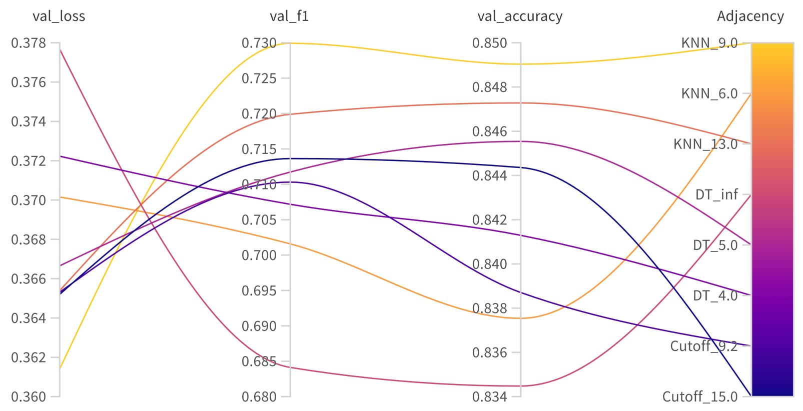


**Fig. S1** Grid sweeping is done with 15 seeds and grouped with adjacency type for $H_{KF}$

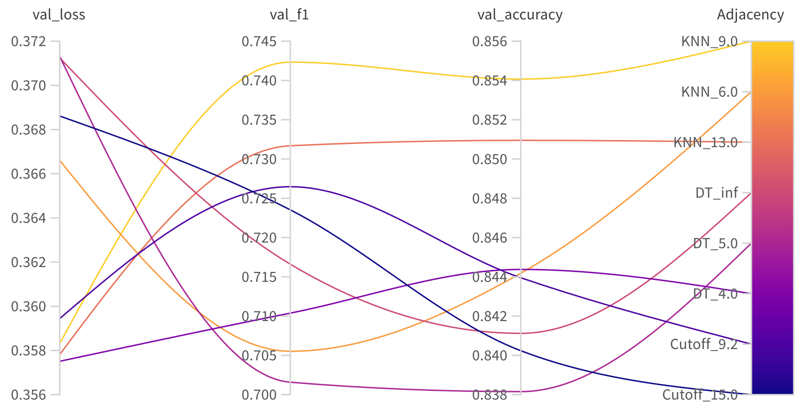


**Fig. S2** Grid sweeping is done with 15 seeds and grouped with adjacency type for $H_{OHE}$

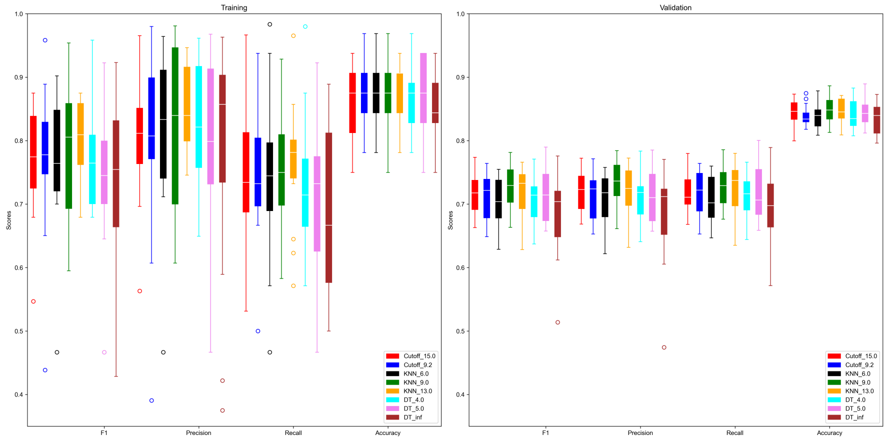


**Fig. S3** Boxplot showing the train and validation score distributions across 15 seeds for the $H_{KF}$ scheme model. We see that most of the distributions are overlapping and comparable.

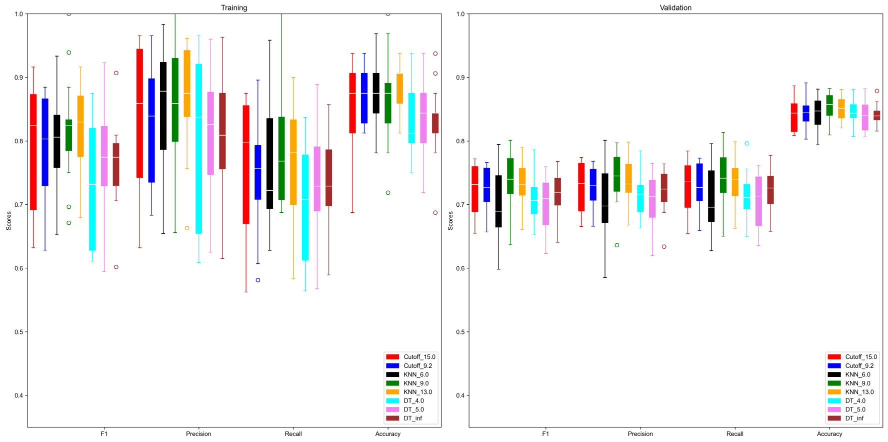


**Fig. S4** Boxplot showing the train and validation score distributions across 15 seeds for the $H_{OHE}$ scheme model. We see that most of the distributions are overlapping and comparable.

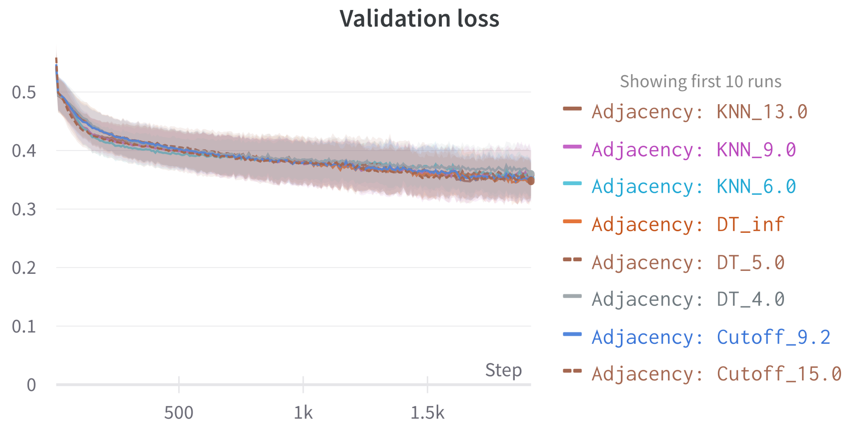


**Fig. S5** Steady/gradual validation step loss decrease with 0.0001 learning rate for the binary classification problem with $H_{KF}$.

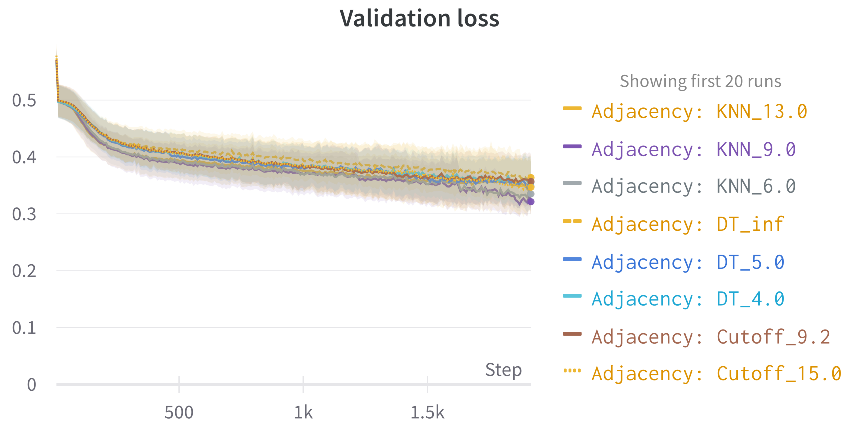


**Fig. S6** Steady/gradual validation step loss decrease with 0.0001 learning rate for the binary classification problem with $H_{OHE}$.

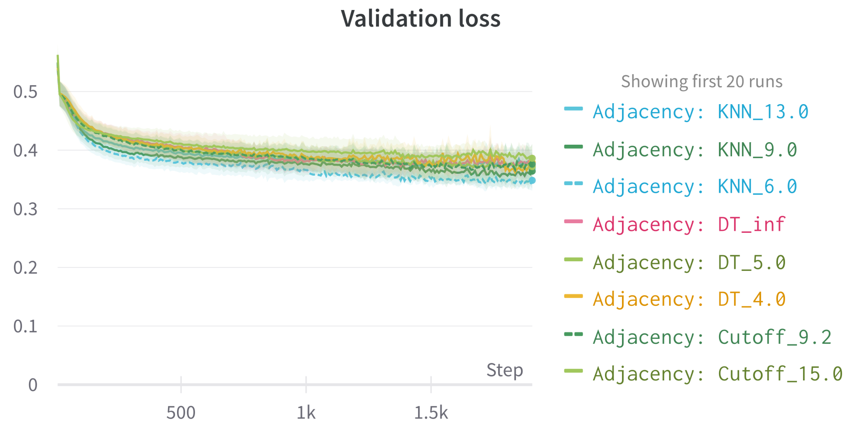


**Fig. S7** Steady/gradual validation step loss decrease with 0.0001 learning rate for the binary classification problem for predicted structures ($H_{KF}$).

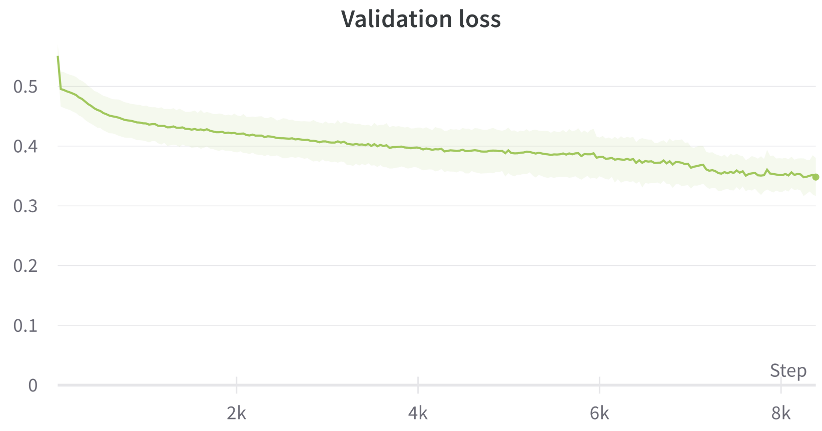


**Fig. S8** Steady/gradual validation step loss decrease with 0.0001 learning rate for the binary classification problem with sequence-based antibody archetype model (DT 5, $H_{KF}$).

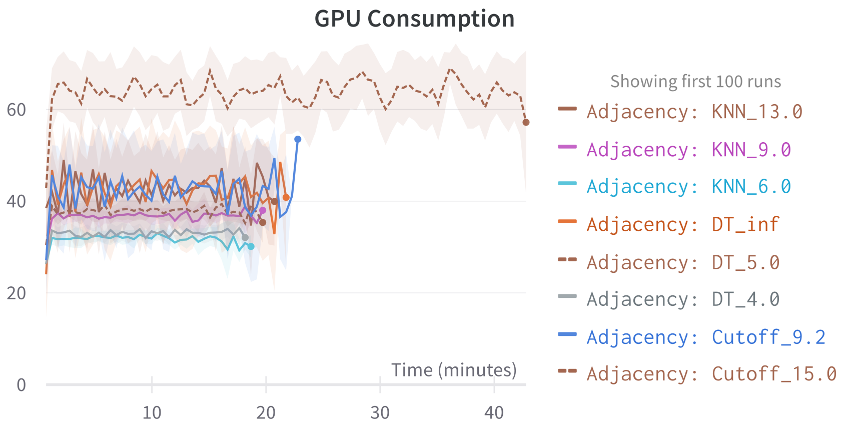


**Fig. S9** GPU consumption for $H_{KF}$

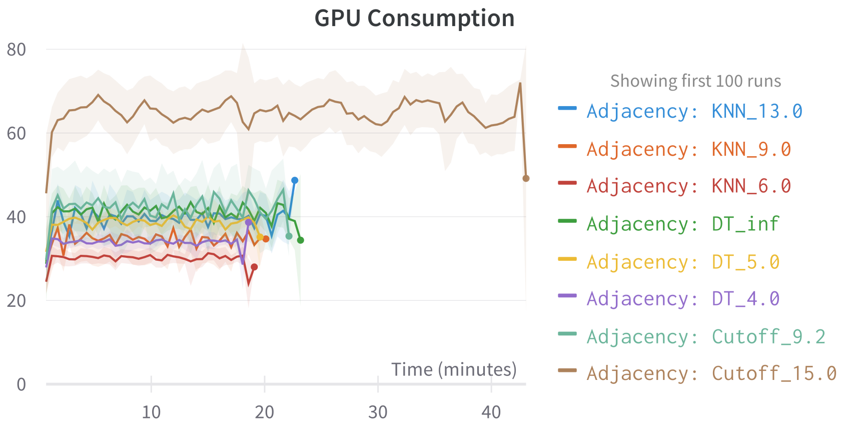


**Fig. S10** GPU consumption for $H_{OHE}$
